# EcRBPome: a comprehensive database of all known *E. coli* RNA-binding proteins

**DOI:** 10.1101/268441

**Authors:** Pritha Ghosh, Niang Guita, Bernard Offmann, R. Sowdhamini

## Abstract

**Background:** The repertoire of RNA-binding proteins (RBPs) in bacteria play crucial role for their survival, and interactions with the host machinery, but there is little information, record or characterisation in bacterial genomes. As a first step towards this, we have chosen the bacterial model system *Escherichia coli*, and organised all RBPs in this organism into a comprehensive database named EcRBPome.

**Results:** EcRBPome contains RBPs recorded from 166 complete *E. coli* proteomes available in the RefSeq database (as of May 2016). The database provides various features related to the *E. coli* RBPs, like their domain architectures, PDB structures, GO and EC annotations etc. It provides the assembly, bioproject and biosample details of each strain, as well as cross-strain comparison of occurrences of various RNA-binding domains (RBDs). The percentage of RBPs, the abundance of the various RBDs harboured by each strain have been graphically represented in this database and available alongside other files for user download.

**Conclusion:** To the best of our knowledge, this is the first database of its kind and we hope that it will be of great use to the biological community. Database URL: http://caps.ncbs.res.in/ecrbpome

## Background

RNA-binding proteins (RBPs) are important regulators of cellular function, being involved in processes at the transcriptional, post-transcriptional, translational, as well as post-translational levels. They mediate transport, stabilisation, metabolism and degradation of transcripts within the cell [1]. Hence, a proper understanding of the ‘RBPome’ of an organism is essential.

The complete RBP repertoire of a few model organisms have now been identified by various research groups, including ours [2–5], but the data is not conveniently available to the users due to the lack of proper organisation. The most widely used of the RBP repositories, RBPDB [6], reports experimentally observed RNA-binding sites that have been manually curated from literature, but was last updated in 2012. This database houses information from *H. sapiens*, *M. musculus*, *D. melanogaster* and *C. elegans*, but not from *E.* coli. The ATtRACT database [7], reported in 2016, lists information on 370 RBPs and 1583 consensus RNA-binding motifs, and compiles experimentally validated data from multiple resources, including RBPDB. The latest version (v 3.0) of the sRNATarBase [8] contains more than 750 small RNA (sRNA)-target entries collected from literature and other prediction algorithms.

Here, we report EcRBPome (http://caps.ncbs.res.in/ecrbpome), a comprehensive database of *E. coli* RBPs. The database documents RBPs identified in all complete *E. coli* proteomes (available in the RefSeq database, as of May 2016) by computational sequence search algorithms and methods as described earlier [9]. The data presented in EcRBPome has been cross-referenced to other popular protein annotation resources, and also made available for user download as parsable and graphical representation files. We hope that this database will be of immense importance to the microbial, and in general to the biological community and can be the start point for understanding RBP-mediated regulation in various other lesser studied species.

## Construction and content

### Datasets

All the data presented in this database were obtained from our previous study [9], in which genome-wide survey (GWS) of RBPs were performed for 166 complete *E. coli* proteomes, retrieved from the RefSeq database (May 2016) (please see **Additional File 1** for further details on the search method). The start-points for such search methods, were known sequence and structure signatures of RBPs, organised as structure-centric and sequence-centric family Hidden Markov Models (HMMs) [5]. 8464 putative RBPs could be identified from 166 *E. coli* proteomes studied (**Table 1**). The RefSeq accession numbers, FASTA sequences, domain compositions and cross-references to other databases of these RBPs have been made available for the users in EcRBPome (‘Browse all RBPs in EcRBPome’ under the Browse menu).

**Table 1.**
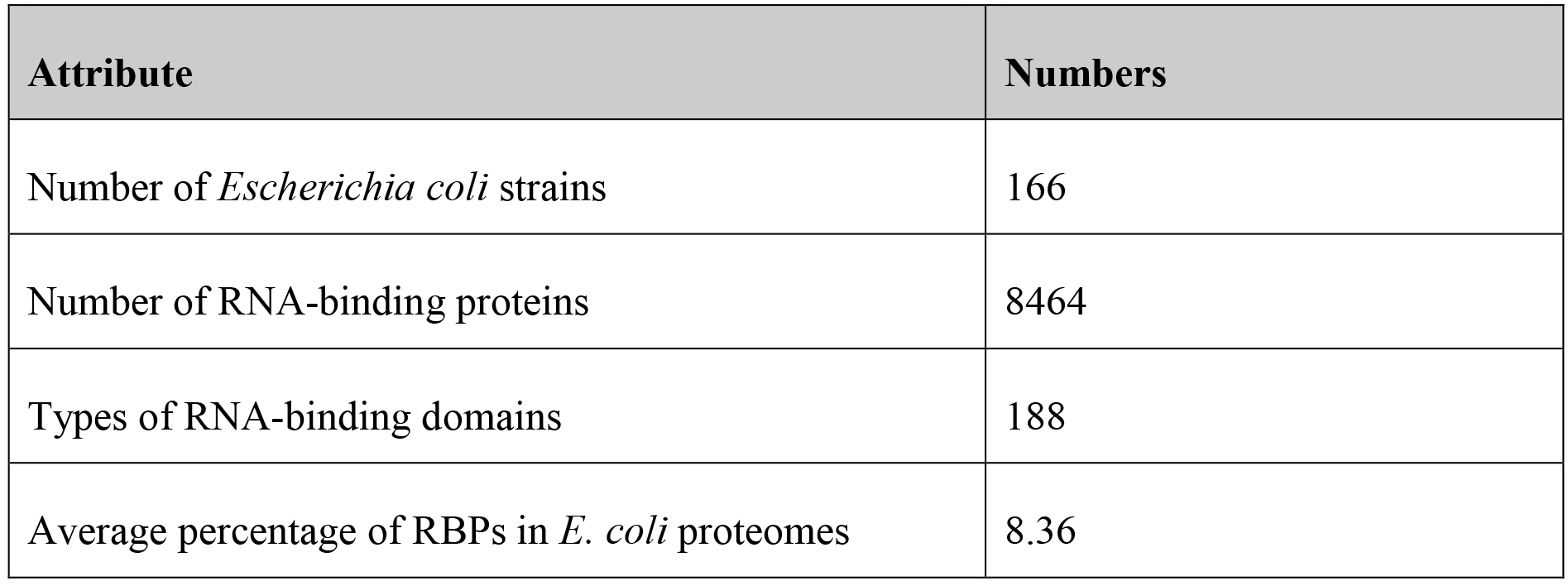
Table of statistics. The various attributes recorded in EcRBPome.

| Attribute | Numbers |
| --- | --- |
| Number of *Escherichia coli* strains | 166 |
| Number of RNA-binding proteins | 8464 |
| Types of RNA-binding domains | 188 |
| Average percentage of RBPs in *E. coli* proteomes | 8.36 |

### Implementation

The retrieval of data and manipulation logic at the back-end of EcRBPome has been implemented using CGI-Perl and the interface of the database built on HTML5, CSS, JavaScript, Ajax and JQuery. The basic tables in EcRBPome have been organised as comma-separated text files, and converted to JSon format, for performance improvement through utilities. The display of tables has been implemented using Bootstrap DataTables. The downloadable graphical plots have been generated using R and the interactive bar plots using the CanvasJS library of JavaScript and HTML5.

### Features

1. **Browse menu:** The users can browse through the list of all the *E. coli* strains present in this database (with links to the assembly, biosample and bioproject details for each strain), all RBPs (with links to the RefSeq page and their downloadable FASTA sequences) and their Pfam28 domain architectures (DAs) [10]. The pathogenic and the non-pathogenic strains have been represented in red and green fonts, respectively. The distribution of various Pfam RBDs and Pfam DAs (domain pairs) in pathogen-specific and nonpathogen-specific proteins have also been represented in various tables (please see **Additional File 1** for more details on the identification of pathogen-specific and nonpathogen-specific proteins). The RBDs, pathogen-specific RBDs and Pfam domain pairs, and nonpathogen-specific RBDs and Pfam domain pairs have been highlighted in bold, red and green fonts, respectively. The sequences of the RBPs can also be submitted to RStrucFam [11], for the prediction of their function and cognate RNA partner(s). **Figure 1A** demonstrates sequence submission to RStrucFam (from the ‘Browse all RBPs in EcRBPome’ option, under the ‘Browse’ menu), followed by the display of results, and navigation to the RStrucFam web server for the details of the identified family(ies).
2. **Cross-strain comparisons**: The various *E. coli* strains present in this database are compared on the basis of different parameters like, percentage of RBPs in each proteome (downloadable graphical representations, as well as comparative account with the average RBP percentage across all strains) (**Figure 1B**), presence or absence of RBDs in each strain (matrix representation) (**Figure 1C**), as well as percentage of the various RBDs in each strain (graphical representations and downloadable tab separated text files) (**Figure 1D**). The RBPs obtained from 166 different *E. coli* strains were compared in terms of sequence, on the basis of single-link clustering method (please see **Additional File 1** for a description of the method). Each of the proteins reported in this database were searched against all eukaryotic proteins present in the UniProt database, using the PSI-BLAST module of the BLAST 2.2.30+ suite [12] (sequence E-value cut-off = 10^−5^, inclusion threshold = 10^−5^, number of iterations = 5). The hits obtained were filtered on the basis of 30% sequence identity and 70% query coverage. The PSI-BLAST results for proteins belonging to each cluster have been made available for download by the users.
3. **Cross-reference to other databases:** EcRBPome provides annotations for each RBP by establishing links to other resources like, UniProt [13] (sequence annotation database), Protein Data Bank (PDB) [14] (structure annotation database) and Gene Ontology (GO)
[15] and Enzyme Commissions (functional annotation resources).
4. **Download sequences:** FASTA sequences of RBPs encoded in each strain, all RBPs present in this database and those of RBDs predicted to be encoded in these RBPs are available for download by the users.

**Figure 1.**
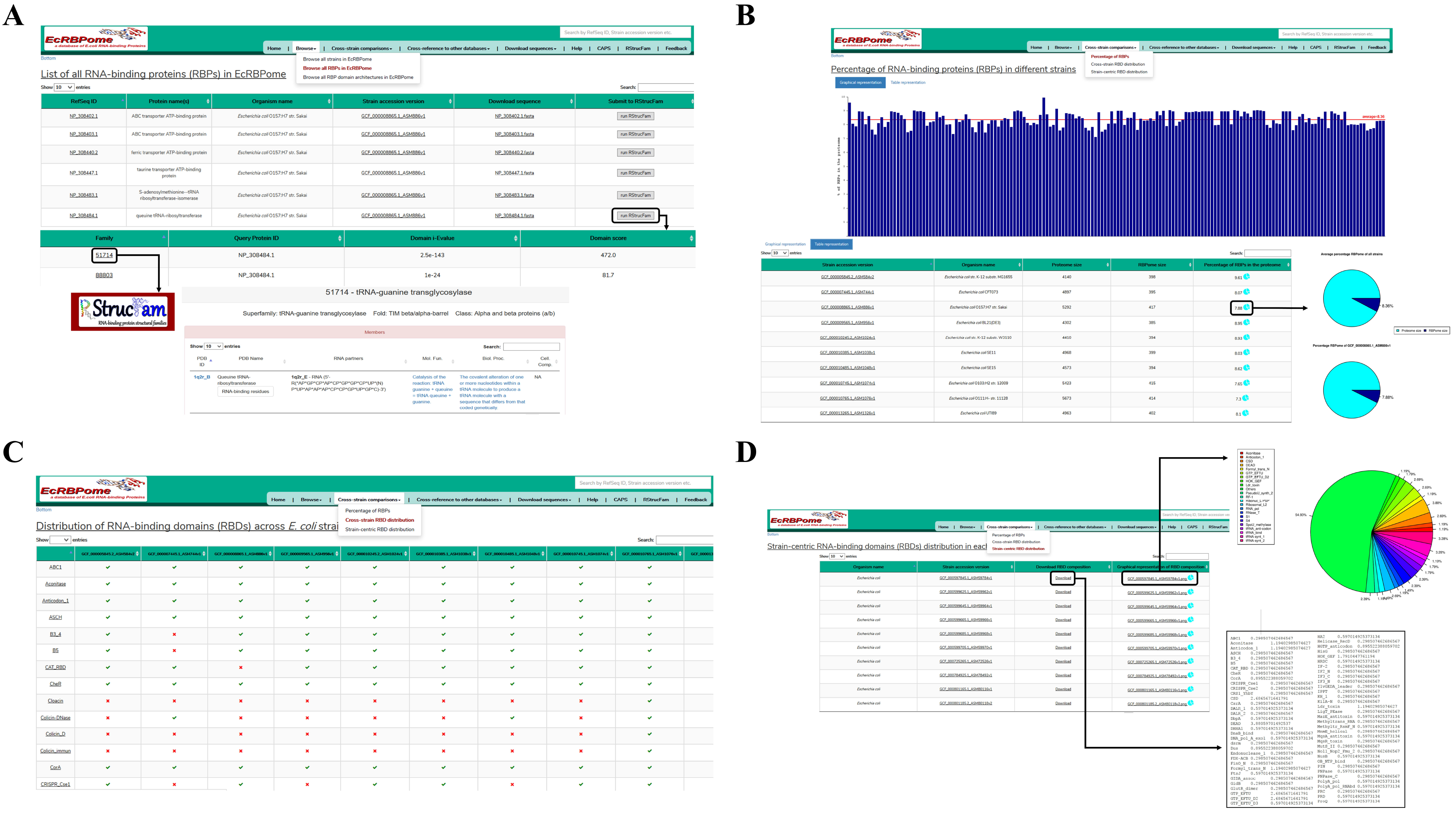
Database organisation and features. The organisation of the EcRBPome database and its important features have been represented in this figure. **A.** Sequence submission to RStrucFam, for the prediction of putative function(s) and cognate RNA partners. The snippets show the results page and the navigation to the RStrucFam web server for the details of the identified family(ies) have also been depicted. **B.** Graphical and tabular representations of the percentage of RBPs in the strains present in this database. Comparative pie-charts for these values in each strain and the average across all strains, are available for user download. **C.** Matrix representations for the distributions of various RBDs across the different *E. coli* strains. Presence of a particular RBD in a strain is denoted with a green tick mark, whereas absence is denoted by a red cross mark. **D.** RBD composition of each strain are available as user downloadable pie charts, as well as tab separated text files.

Further details of the features have been made available in the database ‘Help’ page and also in **Supplementary video**.

## Utility and discussion

To the best of our knowledge, EcRBPome is the first database of its kind that organises all RBPs known in a model organism in one platform. EcRBPome records information from all known complete *E. coli* proteomes (as of May 2016), and also links the data present in this database to other sequence, structure and function annotation resources. Hence, it is a ‘one-stop solution’ for all researchers who prefer to understand the global landscape of *E. coli* RBPs, as well as those who are interested in specific strains or proteins. It also predicts the function(s) and cognate RNA partner(s) for each of the RBPs present in this database, through our in-house algorithm, named RStrucFam. A case study for the same has been discussed in the following section.

### Case study

The ‘MULTISPECIES: riboflavin biosynthesis protein RibD’ (alternately named as ‘MULTISPECIES: bifunctional diaminohydroxyphosphoribosylaminopyrimidine deaminase/5-amino-6-(5 phosphoribosylamino) uracil reductase’) (RefSeq ID: WP_001150457.1), was found to be present in 51 out of the 166 strains recorded in this database. The protein associates with two UniProt entries (IDs: P25539 and Q3ZUB0), and three PDB structures (codes: 2G6V, 2O7P and 2OBC). This protein will be henceforth referred to as ‘query’. The query sequence is predicted to associate with a ‘populated SCOP family’ (ID: 89800) in RStrucFam, which is single-membered (PDB chain ID: 2B3JD; RNA partner chain IDs: 2B3JE, 2B3JF and 2B3JH). Hence, the query protein is also predicted to bind to these aforementioned RNA chains, which are redundant in terms of sequence.

Structural alignment of 2B3JD and largest of the query protein structures, 2G6VA (with the best resolution) were performed using the structural alignment tool, Matt [16]. The RNA-interacting residues in 2B3JD, as predicted by the RStrucFam algorithm, using 5Å distance cut-off criterion, have been highlighted in yellow in **Figure 2A**. The residues in 2G6VA that are structurally aligned with the above-mentioned residues, have been highlighted in cyan in **Figure 2A**, and were used to guide the docking of the RNA chain (2B3JH) onto the protein chain (2G6VA), using the docking tool HADDOCK [17]. The structures of the RNA-protein complexes (2B3JD-2B3JH and 2G6VA-2B3JH) have been shown on the left panes of **Figure 2B** and **Figure 2C**, respectively. The colour coding used to highlight the residues are same as those followed in Figure 2A. The gaps seen on the structures of the proteins and the RNA are due to the absence of ATOM coordinates in those regions of the molecules.

**Figure 2.**
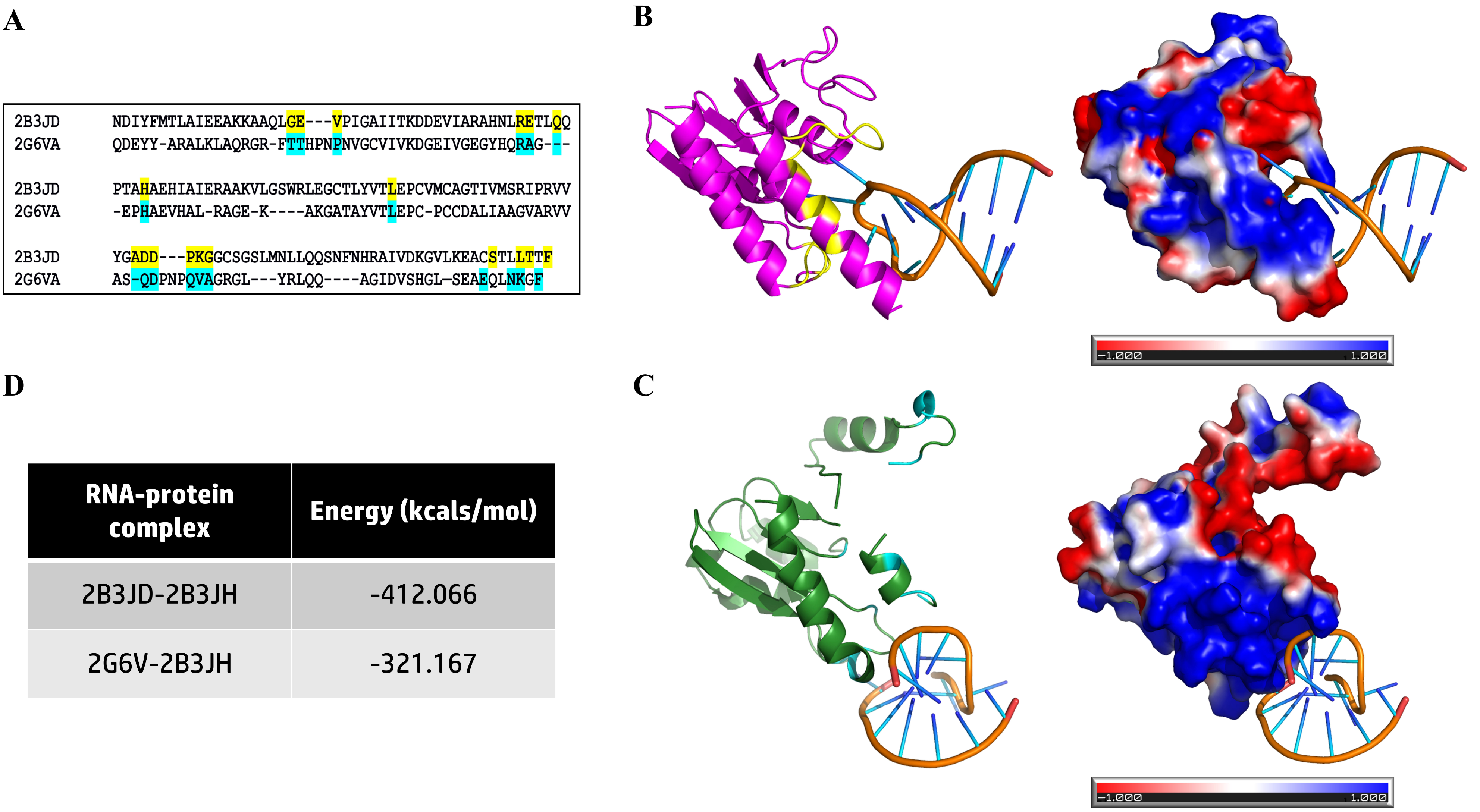
Comparison of RNA-binding affinities of two proteins. The RNA-binding properties of two proteins have been compared in this case study, on the basis of predictions made by RStrucFam. **A.** Structural alignment of the two proteins. The RNA-binding residues in 2B3JD (on the basis of 5Å distance cut-off criterion) have been highlighted in yellow, whereas the structurally aligned residues in 2G6VA have been highlighted in cyan. The same colour scheme have also been followed in panels B and C of this figure. **B.** Structure of the 2B3JD-2B3JH complex (left pane) and its electrostatics properties on the solvent accessible surface (right pane). **C.** Structure of the 2G6VA-2B3JH complex (left pane) and its electrostatics properties on the solvent accessible surface (right pane). **D.** The potential energies of the two complexes (in kcals/mol) have been tabulated. These values were calculated using SYBYL7.2 (Force Field: Tripos, Electrostatics: None) in vacuum, post energy minimisations until convergence.

Electrostatic potential was calculated using PDB2PQR [18] (in the AMBER force field) and Adaptive Poisson-Boltzmann Solver (APBS) [19]. The ±1 *kT*/*e* (where, ‘k’ is the Boltzmann’s constant, ‘T’ is temperature in Kelvin and ‘e’ is the charge of an electron) electrostatic potential on the solvent accessible surfaces of the proteins have been shown on the right panes of Figure 2B and Figure 2C, for the 2B3JD-2B3JH and 2G6VA-2B3JH complexes, respectively. Though the proteins are predicted to belong to the same structural family, their surface electrostatic maps are quite different. It is to be noted that in both the cases, the partner RNA binds amidst a large electropositive patch.

These complexes were subjected to energy minimisations until convergence using SYBYL7.2 (Force Field: Tripos, Electrostatics: None) in vacuum and their potential energy values have been represented in **Figure 2D**. This proves that proteins belonging to the same structural family are capable of binding to the same RNA, but with differential RNA-binding affinities, as seen in our previous studies also [20].

## Conclusions

RBPs and sRNAs play important roles in bacterial post-transcriptional regulation of gene expression, and have been highly studied over the past decade [21,22]. The number of complete genome sequences available has exponentially increased due to the advent of next generation sequencing technologies. Detailed structural and functional characterisation of several RBPs, even within *E. coli* genome, requires pain-staking efforts and huge amounts of time. Computational approaches offer the first glimpse of putative RBPs using mathematical models of known RBPs and searches in whole genomes.

EcRBPome is a comprehensive platform for information on all RBPs from a popular model organism, *E. coli*. Sequences of RBPs reported in this database can also be used to select target gene products for detailed characterisation and to serve as start points for identifying sequence homologues in other microbial proteomes. Especially, the less studied species, where performing studies using experimental techniques are a challenge. For example, gene products of microorganisms that are highly pathogenic or the ones that are difficult to culture in the laboratory could be studied using this approach. The existing study will be further extended to the ever-growing number of complete *E. coli* proteomes and the EcRBPome will be updated with cross-references to a greater number of in-house, as well as external databases and softwares, to enrich the existing repository of information. RBPs can then be followed over taxonomic lineages to understand their patterns of conservation.

## Additional files

**Additional File 1 | Supplementary text**. The text elaborates on the details of the search and the clustering methods used in this study.

**Additional File 2 | Video tutorial of the database**. The video demonstrates the various menus, browse options and other features available in EcRBPome. A text help for the same is available on the database ‘Help’ page.

## Declarations

All authors have gone through the manuscript and contents of this article have not been published elsewhere.

## Ethics

Not applicable, since this study has not directly used samples collected from humans, plant or animals, but has analysed publicly available, pre-existing protein sequence data.

## Consent to Publish

Not applicable.

## Availability of Data and Materials

All the data related to this work, including accession IDs of proteins, have been presented in the database (URL: http://caps.ncbs.res.in/ecrbpome).

## List of abbreviations

APBS: Adaptive Poisson-Boltzmann Solver
DA: Domain architecture
*E. coli*: *Escherichia coli*
GWS: Genome-wide survey
HMM: Hidden Markov Model
PDB: Protein Data Bank
RBD: RNA-binding domain
RBP: RNA-binding protein
sRNA: Small RNA

## Authors’ contributions

RS conceived the idea and designed the project. PG and NG acquired data and performed all the analyses. PG wrote the first draft of the manuscript and RS improved on it. All the authors read and approved the final version of the manuscript.

## Acknowledgments

We thank NCBS (TIFR) for financial and infrastructural support.

